# Transcriptomic analysis of longitudinal *Burkholderia pseudomallei* infecting the cystic fibrosis lung

**DOI:** 10.1101/229823

**Authors:** Erin P. Price, Linda T. Viberg, Timothy J. Kidd, Scott C. Bell, Bart J. Currie, Derek S. Sarovich

## Abstract

The melioidosis bacterium, *Burkholderia pseudomallei*, is increasingly being recognized as a pathogen in patients with cystic fibrosis (CF). We have recently catalogued genome-wide variation of paired, isogenic *B. pseudomallei* isolates from seven Australasian CF cases, which were collected between four and 55 months apart. Here, we extend this investigation by documenting the transcriptomic changes in *B. pseudomallei* in five cases. Following growth in an artificial CF sputum medium, four of the five paired isolates exhibited significant differential gene expression (DE) that affected between 32 and 792 genes. The greatest number of DE events was observed between patient CF9 strains, consistent with the hypermutator status of the latter strain, which is deficient in the DNA mismatch repair protein MutS. Two patient isolates harbored duplications that concomitantly increased expression of the β-lactamase gene *penA*, and a 35kb deletion in another abolished expression of 29 genes. Convergent expression profiles in the chronically-adapted isolates identified two significantly downregulated and 17 significantly upregulated loci, including the antibiotic resistance-nodulation-division (RND) efflux pump BpeEF-OprC, the quorum-sensing *hhqABCDE* operon, and a cyanide- and pyocyanin-insensitive cytochrome *bd* quinol oxidase. These convergent pathoadaptations lead to increased expression of pathways that may suppress competing bacterial and fungal pathogens and that enhance survival in oxygen-restricted environments, the latter of which may render conventional antibiotics less effective *in vivo*. Treating chronically-adapted *B. pseudomallei* infections with antibiotics designed to target anaerobic infections, such as the nitroimidazole class of antibiotics, may significantly improve pathogen eradication attempts by exploiting this Achilles heel.

## INTRODUCTION

The Gram-negative soil-dwelling bacterium *Burkholderia pseudomallei* causes melioidosis, an opportunistic tropical infectious disease of humans and animals that has a high fatality rate (Wiersinga et al. 2012). *B. pseudomallei* is found in many tropical and subtropical regions globally, and has been unmasked in temperate and even arid environments following unusually wet weather events (Yip et al. 2015; Chapple et al. 2016; Sarovich et al. 2016). Infection occurs following percutaneous inoculation from contaminated soil or water, inhalation, or ingestion. Melioidosis symptoms vary widely due to the ability for *B. pseudomallei* to infect almost any organ, with pneumonia being the most common presentation (Leelarasamee and Bovornkitti 1989; Currie et al. 2010). Individuals most at risk of contracting melioidosis include diabetics, those with hazardous alcohol consumption, and the immunosuppressed. There has been increasing recognition that people with chronic lung diseases such as cystic fibrosis (CF) are also at a heightened risk (Holland et al. 2002; O'Carroll et al. 2003).

CF is a heritable disorder of the *CFTR* gene, and defects in CFTR lead to exaggerated and ineffective airway inflammation, an imbalance in salt regulation in the lungs and pancreas, and a chronic overproduction of thick and sticky mucus in the airways and digestive system (Amaral 2015). Impaired immunity and mucus clearance encourage infection and subsequent persistence and adaptation of opportunistic bacterial pathogens in the CF lung, leading to the development of bronchiectasis with subsequent progressive pulmonary decline, and ultimately, loss of pulmonary function and death (Cohen and Prince 2012).

The most common pathogens of the CF lung comprise *Pseudomonas aeruginosa, Staphylococcus aureus, Haemophilus influenzae*, and less commonly, *Achromobacter xylosoxidans*, non-tuberculosis mycobacteria, *Stenotrophomonas maltophilia* and certain *Burkholderia* species, including *B. cepacia* complex species and *B. pseudomallei* (Coutinho et al. 2008). The most common and best-studied CF pathogen is *P. aeruginosa*, which can adapt to the CF lung environment via various mechanisms. Convergent pathoadaptations in *P. aeruginosa* include the downregulation or loss of virulence factors and motility-encoding loci, emergence of hypermutators, enhanced antibiotic resistance and immune evasion facilitated by a switch to mucoidy and a biofilm-based lifestyle, and altered expression of other loci enhancing bacterial metabolism and survival within the nutrient-poor CF lung environment (Cohen and Prince 2012; Winstanley et al. 2016).

Improving life expectancy for those with CF has led to an increased risk of exposure to *B. pseudomallei* following travel to melioidosis-endemic regions. Although uncommon, infection of the CF lung by *B. pseudomallei* has now been documented in at least 25 cases worldwide (Geake et al. 2015). Due to low total case numbers, comparatively little is understood about the pathogenic role of *B. pseudomallei* in CF pulmonary disease. The most common clinical presentation is chronic carriage (76%), which is associated with accelerated lung function decline (Geake et al. 2015). This prevalence contrasts with melioidosis in non-CF patients, where chronic carriage is exceedingly rare, occurring in only 0.2% of cases (Price et al. 2015). To better understand *B. pseudomallei* pathoadaptation in the CF lung, we recently investigated the genome-wide evolution of isogenic *B. pseudomallei* strains isolated from seven Australasian CF patients, which were collected between 4 and 55 months apart (Viberg et al. 2017s). Hallmarks of these infections included *B. pseudomallei* persistence despite multiple eradication attempts, multidrug resistance, mutations in virulence, metabolism and cell wall components, and the first-documented case of hypermutation in *B. pseudomallei*. In all except one case, multiple single-nucleotide polymorphism (SNP) and insertion-deletion (indel) mutations were identified, with a high rate of nonsynonymous mutations, many of which were predicted to affect protein function (Viberg et al. 2017).

RNA-seq provides a detailed view of the transcriptional landscape in bacterial isolates grown under different conditions or niches (Sharma et al. 2010), and is now a well-established method for examining differential gene expression (DE) in bacterial pathogens (Creecy and Conway 2015) . Here, we performed bacterial RNA-seq on five of the CF cases that we have recently described (Viberg et al. 2017) to catalogue both within-host and convergent transcriptional evolution during long-term *B. pseudomallei* infection in the CF lung. Paired isolates representing the initial and the most recent cultures available from each patient were compared. *B. pseudomallei* cultures were grown in an artificial sputum medium (Sriramulu et al. 2005; Fung et al. 2010) to mimic the conditions found in the CF lung environment.

## RESULTS

### Differential gene expression among CF isogenic pairs

DE was observed in four of the five CF pairs, with only the CF10 pair failing to yield significant transcriptional differences (i.e. no genes with a ≥1.5 log_2_-fold change and a false discovery rate (FDR) of ≤0.01; Figure 1). We have previously shown that no genetic variants separate the CF10 strains, which had the shortest time between collection of only 10 months (Viberg et al. 2017). This lack of significant transcriptional differences rules out epigenetic effects on gene expression between this pair, at least under the tested growth conditions, and illustrates that RNA-seq is a robust methodology that is not readily prone to false-positive results.

**Figure 1.**
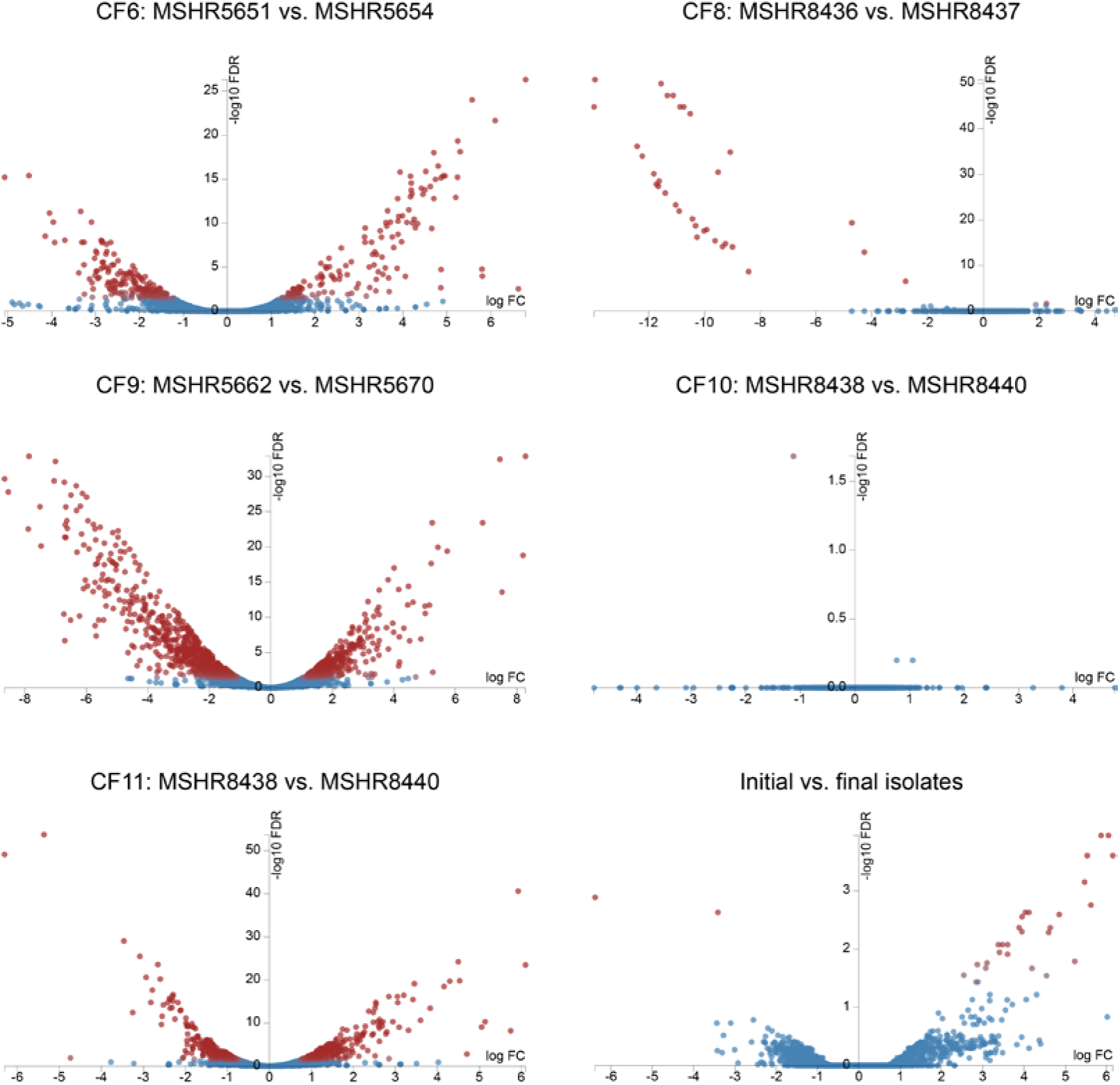
Degust volcano plots showing differentially expressed (DE) genes between paired *Burkholderia pseudomallei* isolates retrieved from five cystic fibrosis (CF) lungs, and between initial and latter isolates. Four of the five pairs exhibited DE; CF10, with the shortest time between isolates, exhibited no genetic or significant transcriptomic changes. CF6, CF8, CF9 and CF11 pairs were separated by 229, 32, 792 and 169 DE loci, respectively. The 32 DE loci in CF8 were all downregulated; DE genes in CF6, CF9 and CF11 were down- or up-regulated. Nineteen loci were DE between initial and latter isolates, of which 17 were upregulated. Blue, genes with no significant DE; red, genes with significant DE.

Of the four pairs with significant DE, the CF8 pair had the least with 32 loci, followed by CF11, CF6 and CF9 with 169, 229 and 792 DE loci, respectively (Figure 1; Table 1). These paired isolates were collected 46, 14, 27 and 55 months apart, respectively. There was good correlation between the proportion of DE loci and the genome-wide mutations catalogued between these isolate pairs (Viberg et al. 2017), with 12, 15, 24, and 112 mutational events (i.e. SNPs, indels, deletions or gene duplications) identified in CF8, CF11, CF6 and CF9, respectively (Table 1). The elevated number of mutations seen in CF9 is due to a *mutS* mutation in the latter strain, which confers a hypermutator phenotype, the first time hypermutation has been described in *B. pseudomallei* (Viberg et al. 2017); this in turn contributes to a high number of DE genes. However, when comparing the ratio of DE genes to mutational events, CF11 had the highest proportion of DE genes (11.3), followed by CF6 (9.5), CF9 (7.1) and CF8 (2.7). There was a significant skew in DE towards genes located on chromosome II, which contains a lower proportion of housekeeping genes than chromosome I (Holden et al. 2004). Despite encoding only 44% of the genome by size and 41% of coding sequences, chromosome II loci were significantly overrepresented in the non-hypermutator CF pairs (Pearson’s X^2^ test *p*<0.001), with between 63 and 97% of the DE genes residing on chromosome II. In CF9, there was a non-significant trend towards chromosome II loci, with 52% of DE loci located on this chromosome, pointing to the more random nature of mutations in the CF9 hypermutator compared with the other cases (Table S1).

**Table 1.** Summary of the genetic mutations and differentially expressed (DE) genes between paired, sequential *Burkholderia pseudomallei* isolates obtained from five cystic fibrosis patients^a^.

| Patient | Initial and final isolate IDs | Months between collection | No. genome-wide mutational events (Chr I, Chr II) | No. genes affected by mutations (Chr I, Chr II) | No. DE genes (Chr I, Chr II) | No. DE genes (downregulated, upregulated) |
| --- | --- | --- | --- | --- | --- | --- |
| CF6 | MSHR5651, MSHR5654 | 27 | 24 (14, 10) | 29 (14, 15) | 229 (69, 160) | 229 (124, 105<sup>b</sup>) |
| CF8 | MSHR8436, MSHR8437 | 46 | 12 (7, 5) | 39 (5, 34) | 32 (1, 31) | 32 (32<sup>c</sup>, 0) |
| CF9 | MSHR5662, MSHR5670 | 55 | 112 | 79 (40, 39) | 792 (381, 411) | 792 (558, 234) |
| CF10 | MSHR8438, MSHR8440 | 10 | 0 | 0 | 0 | 0 |
| CF11 | MSHR8441, MSHR8442 | 14 | 15 | 38 (10, 28) | 169 (62, 107) | 169 (68<sup>d</sup>, 101) |

<sup>a</sup>Adapted from (Viberg et al. 2017).
<sup>b</sup>Seven of these genes are upregulated due to a 30x duplication affecting these loci in MSHR5654.
<sup>c</sup>Twenty-nine of these genes are downregulated due to a 35kb deletion affecting these loci in MSHR8437.
<sup>d</sup>Twenty-five of these genes appear as downregulated due to a 10x duplication affecting a 36.7kb locus in MSHR8441 compared with only a 2x duplication in MSHR8442 (Viberg et al. 2017).

### Many DE genes are absent in *Burkholderia mallei* and in chronic-carriage melioidosis Patient 314

*B. mallei*, the causative agent of glanders, is an equine-adapted clone of *B. pseudomallei* that continues to undergo dramatic reductive evolution, having already shed ~1.3-1. 5Mbp of its genome since its divergence from *B. pseudomallei* (Holden et al. 2004; Price et al. 2013). As a consequence, this bacterium does not survive in the environment, although it remains highly pathogenic (Dvorak and Spickler 2008). *B. mallei* therefore provides a useful comparison for examining DE in the CF *B. pseudomallei* strains due to similar genome-wide patterns of locus loss and adaptation to a mammalian host (Viberg et al. 2017). We also compared the DE loci in the CF strains to those genes mutated in chronic-carriage melioidosis case P314. P314 has the longest *B. pseudomallei* infection ever documented, and despite multiple eradication attempts, continues to harbor this bacterium in her lungs since she was first diagnosed in 2000. The genome of a 139-month isolate from P314, MSHR6686, shows dramatic adaptation to the lung environment, including the loss of 285kb of chromosome II at four separate locations that collectively encompass 221 genes; ~50% of these genes are also absent in *B. mallei* (Price et al. 2013).

When compared with genes lost in 17 *B. mallei* strains (092700E 11, 2000031063, 2000031281, 2002721280, 6, A188, A193, ATCC10399, ATCC23344, BMQ, China5, China7, FMH23344, GB8horse4, JHU, and PRL-20), there was a 23, 27, 33 and 100% overlap with DE genes in the latter isolates from CF9, CF6, CF11 and CF8, respectively, and when compared with genes lost in MSHR6686, there was a 9, 15, 20 and 97% overlap, respectively (Table S1). The proportion of downregulated genes varied across this dataset, ranging from 13% for CF11 to 100% for CF8, demonstrating that the effect on gene expression at these loci is not unidirectional, with certain overlap loci in fact being upregulated in CF6, CF9 and CF11. Of note, all 29 genes (*BPSS1131-BPSS1159*) that were lost due to a large deletion in the latter CF8 isolate (Viberg et al. 2017) were also absent in *B. mallei* and MSHR6686 (Price et al. 2013), providing further evidence of their dispensability for long-term *B. pseudomallei* survival in the mammalian host. As expected, DE showed dramatic downregulation of between 8- and 14- log2 fold (341 to 15,886x) of these loci (Figure 1).

### DE of surface antigens in the CF6, CF9 and CF11 pairs

*B. pseudomallei* produces four capsular polysaccharides (CPS-I to -IV) and a lipopolysaccharide (LPS). However, only CPS-I (encoded by *BPSL2786-BPSL2810*) and LPS (encoded by *BPSL1936* and *BPSL2672-BPSL2688*) are associated with virulence in mammals (Reckseidler-Zenteno et al. 2010; Stone et al. 2014). The remaining three capsule clusters are found in *B. thailandensis* but are not present or intact in *B. mallei*. Our previous genomic analysis of the CF pairs identified missense mutations affecting the LPS loci *wzt* (*BPSL2681*) and *rmlA* (*BPSL2685*) in CF6, a missense mutation affecting the putative LPS biosynthesis gene *BPSL1119* in CF11, and a CPS-I frameshift mutation affecting *wcbA* (*BPSL2809*) in CF11 (Viberg et al. 2017). Consistent with being under heavy selection during chronic infection, we observed DE of several surface polysaccharide loci in the CF6, CF9 and CF11 pairs, the most dramatic of these being in CF9, with forty-six downregulated surface polysaccharide genes (Table S1).

The LPS loci *wbiE* (*BPSL2676*) and *wbiD* (*BPSL2677*) were downregulated by ~1.8-fold (3x) in the latter CF9 isolate. In addition, the poorly characterized LPS biosynthesis-related membrane protein loci *BPSS1683-BPSS1685* were downregulated by ~5.9-fold (60x). This isolate also exhibited downregulation of all CPS-I loci (except *wcbC*) by between 1.7- and 3.1-fold (3 to 8x). In contrast, the latter isolate from CF11 showed upregulation of the CPS-I loci *BPSL2793-BPSL2797* (*wcbN-wcbM-gmhA-wcbL-wcbK*) when compared with its initial isolate, with increases ranging from 2.7 to 6.1-fold (7 to 69x). However, when compared with initial isolates from CF6, CF8, CF9 and CF10, expression of *BPSL2793*-*BPSL2797* in the latter CF11 strain was in fact downregulated (3.4 to 4.5-fold; 11 to 23x). This observation was confirmed as significant downregulation of these loci in the initial CF11 strain (by between 6.1- and 10.1-fold; 69 to 1,098x) when compared with all other initial strains, rather than significant upregulation of these CPS-I loci in the latter CF11 strain.

Unlike CPS-I, expression of the CPS-II cluster (*BPSS0417-BPSS0429*) is induced when grown in water, suggesting that this polysaccharide plays a role in environmental survival (ReckseidlerZenteno et al. 2010). One locus involved in CPS-II biosynthesis, *BPSS0425*, was downregulated (1.8-fold; 4x) in CF6, and the entire cluster was downregulated in CF9 (range: 2.5- to 5.2-fold; 6 to 37x). Conversely, *BPSS0417* and *BPSS0418* were upregulated in CF11 (1.6- and 1.7-fold; 3x), respectively. However, as with CPS-I, both CF11 strains exhibited significant downregulation of *BPSS0417* and *BPSS0418* when compared with initial strains from CF6, CF8, CF9 and CF10 (2.0- and 3.1-fold; 4 to 9x). The genes encoding CPS-III (*BPSS1825-BPSS1835*) were significantly downregulated in the latter isolates from CF6 (2.4- to 3.1-fold; 5 to 9x) and CF9 (5.6- to 7.5-fold; 50 to 178x). Finally, two genes within the CPS-IV cluster (*BPSL2769-BPSL2785*) were downregulated in CF9 (*BPSL2782* and *BPSL2785;* 1.8- to 2.8- fold; 3 to 7x) but 11/17 loci from this cluster were upregulated in CF11 by 1.7- to 5.9-fold (*BPSL2769, BPSL2775-BPSL2784*). However, unlike CPS-I and CPS-II, this pattern of upregulation in both CF11 isolates was maintained for CPS-IV loci *BPSL2769* and *BPSL2775-BPSL2781* even when compared with the initial CF6, CF8, CF9 and CF10 isolates (2.0- to 4.0- fold; 4 to 16x).

### DE of other virulence-associated loci

In addition to CPS-I and LPS, *B. pseudomallei* encodes for several other virulence factors that enhance organism survival and replication upon infection or that subvert or disarm host defenses. These factors include adhesins, flagella, fimbriae, pili, specialized secretion systems, actin motility proteins, secreted factors and secondary metabolites (Stone et al. 2014). Although virulence factors are often critically important during the acute stages of infection, they can become disadvantageous for long-term survival, presumably due their immunogenicity (Price et al. 2013; Winstanley et al. 2016). Consistent with loss-of-virulence as a pathoadaptive mechanism in chronic *B. pseudomallei* infections, we have previously documented missense mutations affecting the Type 3 secretion system 3 (T3SS-S) gene *bsaW* in CF6, and *Burkholderia* biofilm factor A *bbfA* and fimbrial protein BPSL1628 in CF9 (Viberg et al. 2017). When examining RNA-seq profiles, other virulence genes lacking genetic mutations were found to be significantly downregulated in the latter CF isolates. These loci included three Type IV pilus 7 (TFP7) loci (*pilR, pilG* and *pilN*; 1.7-fold; 3x), the lysozyme inhibitor *BPSL1057* (3.4-fold; 11x), Burkholderia lethal factor 1 (3.2-fold; 10x), four T3SS-3 loci (*bsaS, bsaP, bsaO* and *bsaN*), and 16 flagellum loci in CF9 (average 2.5-fold; 6x), and a trimeric autotransporter adhesin (*bpaC; BPSL1631*) in CF11 (1.6-fold; 3x). Of these, the four T3SS-3 loci are also missing in chronic P314 isolates.

### Several regulators with decreased transcription in the CF isolates are absent in *B. mallei*

We hypothesized that downregulation of transcriptional regulators, particularly those absent in *B. mallei*, would be identified in the latter CF isolates due to niche adaptation. As predicted, two of the four pairs exhibited significant downregulation of transcriptional regulators. The first of these, the Fis family regulatory protein YfhA (encoded by *BPSL0350*), was downregulated by ~2.7-fold (~7x) in both CF6 and CF9. In *E. coli*, Fis is a global regulator that is induced under nutrient-rich conditions and plays a role in the regulation of myriad processes including the initiation of DNA replication, ribosomal RNA transcription activation and capsule expression (Beach and Osuna 1998). Thus, the downregulation of Fis in CF6 and CF9 may be responsible for concomitant downregulation of CPS-II and CPS-III loci in these two patients, among other loci. Additional regulators downregulated in CF9 that are absent in *B. mallei* include the transmembrane regulator PrtR (*BPSL0069;* 2.7-fold), two LysrR-family transcriptional regulators (*BPSS0438* and *BPSS2207;* both 1.7-fold) and the metal-related two-component system response regulator IrlR2 (*BPSS1994;* 2.7-fold).

### High-level TMP/SMX resistance in *B. pseudomallei* involves *bpeEF-oprC* upregulation

The combination antibiotic trimethoprim/sulfamethoxazole (TMP/SMX), the antibiotic of choice in the eradication phase of melioidosis treatment (Lipsitz et al. 2012), was administered to CF6, CF9 and CF11 during their *B. pseudomallei* eradication attempts. Acquired resistance towards this antibiotic emerged in the latter isolates from CF6 and CF11, and in midpoint isolates from CF9 (Viberg et al. 2017). The resistance-nodulation-cell division (RND) efflux pump, BpeEF-OprC, is responsible for widespread TMP resistance in *B. pseudomallei* and has been implicated in TMP/SMX resistance (Podnecky et al. 2017). BpeEF-OprC (*BPSS0292-BPSS0294*) expression is under the control of two LysR-type regulators, BpeT (BPSS0290) and BpeS (BPSL0731). We therefore expected to observe upregulation of *bpeEF-oprC* in CF strains with elevated TMP/SMX MICs, consistent with defective *bpeT* or *bpeS* loci.

The latter isolate from CF6, which encodes a T314fs mutation in *bpeT* and is highly resistant towards TMP/SMX (MIC ≥32 μg/mL), showed 5.2 to 6.8-fold upregulation of *bpeEF-oprC* (38 to 111x; Table S1). This isolate also harbours an R20fs mutation in *ptr1* (*BPSS0039; folM*), which encodes a pteridine reductase that is involved in TMP/SMX resistance (Podnecky et al. 2017); this frameshift truncates Ptr1 from 267 to 91 residues. Both strains isolated from CF11 are also highly resistant to TMP/SMX (MIC ≥32 μg/mL) and encode the BpeS missense variants V40I and R247L (K96243 annotation) when compared with wild-type *B. pseudomallei* strains (Viberg et al. 2017). They also encode a three-residue in-frame insertion of R20-A22 in Ptr1 (Podnecky et al. 2017; Viberg et al. 2017). Because both CF11 strains are TMP/SMX-resistant, DE was determined by comparison with TMP/SMX-sensitive isolates in our dataset. Using this approach, significant upregulation was observed for BpeEF-OprC (5.3 to 6.9-fold; 40 to 121x). DE was not observed for *ptr1* or other genes involved in the folate biosynthesis pathway in either the CF6 or CF11 isolates.

### Ceftazidime resistance can occur by upregulation of *penA*

Ceftazidime (CAZ) is a third-generation cephalosporin antibiotic that is the most commonly recommended therapy for the primary phase of melioidosis treatment (Currie 2015). In addition to TMP/SMX, the latter isolate from CF6 is highly resistant to CAZ (MIC ≥256 μg/mL) (Viberg et al. 2017). High-level CAZ resistance is often conferred by a C94Y substitution (C69Y using Ambler’s (Ambler et al. 1991) numbering scheme) in PenA β-lactamase (Sam et al. 2009; Rholl et al. 2011; Sarovich et al. 2012). We have recently shown that the latter CF6 strain also harbors a ~30x duplication of a 7.5kb region that encompasses *penA*; all 30 copies encode the C69Y variant of this enzyme (Viberg et al. 2017). Consistent with this duplication event, *penA* (*BPSS0946*) expression increased by 4.5-fold (22x) in the latter CF6 strain. Six proximal genes (*BPSS0945; BPSS0948-BPSS0952*) were also upregulated by 3.1- to 4.7-fold (9 to 26x; Table S1). One of these, *BPSS0945*, is a peptidase and a putative virulence factor that may play a role in multinucleated giant cell formation (Singh et al. 2013).

A gene duplication event encompassing *penA* has also been documented in the CF11 isolates. The initial strain showed an elevated MIC towards CAZ (12 μg/mL), corresponding with a ~10x duplication of a 36.7kb region that includes *penA*, whereas the latter strain had a 2x duplication of this region and a wild-type CAZ MIC (2 μg/mL) (Viberg et al. 2017). As expected, *penA* was downregulated by 2.1-fold (4x) in the latter isolate due to five times fewer copies of this gene. Downregulation of other genes within the 36.7kb locus ranged from 1.4 to 3.3-fold (3 to 10x; Table S1).

### Increased doxycycline MICs in CF11 are due to *amrAB-oprA* upregulation and *BPSL3085* mutation

The RND efflux pump, AmrAB-OprA (*BPSL1802-BPSL1804*), efficiently effluxes aminoglycoside- and macrolide-class antibiotics (Moore et al. 1999). We have recently shown that synergistic mutations affecting both its regulator AmrR (*BPSL1805*) and an S-adenosyl-L-methionine (SAM)-dependent methyltransferase (*BPSL3085*) led to doxycycline resistance in an Australian melioidosis case (Webb et al. 2017). Doxycycline was administered to CF11 in combination with TMP/SMX as part of a final attempt to eradicate *B. pseudomallei* (Currie 2015; Geake et al. 2015). This lengthy administration led to a doxycycline MIC of 4-8 μg/mL in the CF11 isolates, both of which were retrieved post-treatment (Viberg et al. 2017).

Both CF11 strains encode a large deletion in *amrR* (*amrR*^ΔV62-H223^). This mutation results in a 2.6- to 3.2-fold (6 to 9x) upregulation of *amrAB-oprA* in these isolates. In addition, a previously undocumented S130L mutation in BPSL3085 likely contributes to the decreased susceptibility observed towards doxycycline.

### Evidence of convergent DE between early and latter CF isolates

Finally, we performed a comparison of expression profiles from all CF cases to identify a signal of convergent gene expression (pathoadaptation) across early vs latter isolates. To yield the most robust and relevant analysis, we excluded the latter isolate from CF10 due to a lack of DE in this strain, and the initial isolate from CF11, which was retrieved >3 years after infection and had already undergone substantial genetic and transcriptional modifications. Exclusion of both strains was supported by a lack of convergent signal when they were included in the analysis (data not shown). Using these parameters, 17 genes were found to be significantly upregulated, and two were significantly downregulated (Table S2). Five (26%) loci encode for hypothetical proteins with no known function, of which four were upregulated. Among the convergently upregulated genes with known function was the RND efflux pump BpeEF-OprC (4.8- to 6.1-fold; 28 to 69x), the CydAB cytochrome *bd* quinol oxidase (5.5- to 5.9-fold; 45 to 60x), and the quorum sensing *hhqABCDEFG (BPSS0481-BPSS0487*) operon (3.4- to 4.1-fold; 11 to 17x). The downregulated locus, *BPSS0351*, encodes the MerR family transcriptional regulator CueR (3.4-fold; 11x).

## DISCUSSION

The causative agent of the tropical disease melioidosis, *B. pseudomallei*, is an uncommon pathogen in CF, with fewer than 30 cases documented worldwide to date (Geake et al. 2015). We have recently performed comparative genomic analysis of isogenic strains collected between 4 and 55 months apart from the airways of seven of these cases (Viberg et al. 2017). Here, we sought to further characterize these chronic cases by examining the transcriptomes of five paired *B. pseudomallei* isolates retrieved between 10 and 55 months apart. Isolates were cultured in an artificial CF sputum medium (Fung et al. 2010) to mimic their original *in vivo* environment.

Under these conditions, DE was detected in four of the five cases and ranged from 32 to 792 genes, with the hypermutator strain from CF9 contributing the greatest number of DE loci (Table S1). Interestingly, when compared with the number of genetic changes occurring in each isolate pair, the latter isolate from CF11 had a higher proportion of DE loci to mutations (11.3) than CF9 (7.1), demonstrating that hypermutation does not necessarily lead to a similarly high number of transcriptional differences. The one case with no DE, CF10, exhibited no genetic changes (i.e. SNPs, small indels, copy-number variants, or large deletions) and had the shortest time between isolate collection at 10 months (Viberg et al. 2017). All other cases encoded genetic differences between pairs. The DE genes fell into several functional categories (Table S1), reflecting the diversity and versatility of pathoadaptive pathways in *B. pseudomallei*. Our RNA-seq analysis revealed that many of the DE genes were absent in the chronic P314 strain (range: 9-97%) or in *B. mallei* (range: 23-100%), providing further evidence that these loci are not required for long-term survival in the human airways. Perhaps most striking was the observation that nearly one-third (32%) of DE genes lack a known function, highlighting the relative paucity of functional studies into this important yet under-recognized pathogen.

*B. pseudomallei* has a ~7.3Mbp genome that is encoded on two replicons; a ~4.1Mbp ‘housekeeping’ chromosome I, and a ~3.2Mbp ‘accessory’ chromosome II. The genome of archetypal strain K96243 consists of 3,460 and 2,395 coding sequences on chromosomes I and II, respectively (Holden et al. 2004). There was a greater proportion of DE genes (between 52 and 97%) on chromosome II in all cases, despite its smaller size and fewer coding sequences. This bias towards DE on the accessory chromosome contrasts with a 2009 study of a CF-derived *B. cenocepacia* isolate, which displayed a greater proportion of DE genes on chromosome I compared with chromosomes II and III when the isolate was grown in an artificial sputum medium (Yoder-Himes et al. 2009). However, the study by Yoder-Himes and colleagues compared DE of a single isolate grown in sputum versus a soil medium, rather than between longitudinal clinical isolates, which may explain the discordance between studies. We have previously shown that a greater proportion of mutational events affect chromosome II of *B. pseudomallei* in a long-term chronic-carriage isolate from P314 (Price et al. 2013), and in the mammalian-adapted *B. pseudomallei* clone *B. mallei*, a greater proportion of genes have been lost from chromosome II than chromosome I, with chromosome II representing 44% of the *B. pseudomallei* K96243 genome (Holden et al. 2004) but only 40% of the ATCC 23344 *B. mallei* genome (Nierman et al. 2004). The skew towards DE loci on chromosome II in chronic *B. pseudomallei* isolates points towards a lesser role for chromosome II loci in bacterial survival and persistence within the human host, which is reflected by the greater degree of reductive evolution affecting this replicon (Price et al. 2013; Viberg et al. 2017). In their study of the transcriptional landscape of *B. pseudomallei*, Ooi and co-workers found that only ~28% of chromosome II genes were expressed under a single condition, compared with ~72% of chromosome I genes (Ooi et al. 2013). Taken together, these results confirm the ‘accessory’ role of the chromosome II replicon in *B. pseudomallei*.

Attenuation of immunogenic surface antigens and other virulence factors are hallmarks of chronically persistent infections across many pathogenic bacterial species, including *B. pseudomallei* (Price et al. 2013; Viberg et al. 2017). Encoded by the 34.5kb *wcb* operon *BPSL2786-BPSL2810* (Reckseidler et al. 2001; Reckseidler-Zenteno et al. 2005), the *B. pseudomallei* CPS-I is a potent virulence determinant that imparts high-level serum resistance and facilitates phagocytic evasion (Reckseidler-Zenteno et al. 2005). This capsule is also intact in *B. mallei* and has been shown to be essential for its virulence (DeShazer et al. 2001 ; Atkins et al. 2002). Our prior genomic analysis identified only a single CF pair with mutated CPS-I in CF11. This frameshift mutation in *wcbA* results in a truncated protein (Viberg et al. 2017) that would likely cause reduced, although not abolished, CPS-I production (Cuccui et al. 2012). The DE analysis provides further evidence of CPS-I inactivation in the CF pairs, with downregulation of all but one of the CPS-I loci in the latter CF9 isolate, and downregulation of *wcbN-wcbM-gmhA-wcbL-wcbK* in both CF11 isolates. In both cases, it is likely that CPS-I production was either substantially reduced or abolished. Although CPS-III is not required for virulence, it is noteworthy that this locus was also downregulated in CF9 and CF11, as it has been previously shown that CPS-III expression is tied to that of CPS-I genes (Ooi et al. 2013).

Like CPS-I, the *B. pseudomallei* LPS is required for capsule biosynthesis, virulence and serum resistance (DeShazer et al. 1998). Its immunogenic outer membrane component is readily recognized by the host innate immune system (Tuanyok et al. 2012), which makes LPS a target for inactivation in chronic bacterial infections. We have previously uncovered missense mutations in the LPS *wzt* and *rmlA* loci of CF6, and a missense mutation affecting *BPSL1119* in CF11 (Viberg et al. 2017). DE analysis identified additional evidence for reduced or abolished LPS production in a third CF case, CF9, due to the significant downregulation of *wbiD, wbiE*, and *BPSS1683-BPSS1685*. The convergent evolution of the chronic strains to attenuate CPS-I and LPS loci demonstrates the dispensable, and probably highly unfavourable, nature of these surface antigens for long-term survival of this pathogen in the CF airways. Prior work has suggested that CPS-I and LPS may be disadvantageous for *B. pseudomallei* persistence due to their virulence potential and immunogenicity (Price et al. 2013). By examining both genomic and transcriptomic modifications over time, it is now clear that these capsule clusters, in their wildtype form, pose a major issue for successful long-term *B. pseudomallei* persistence in the CF lung, with the bacterium either mutating or downregulating key genes in CPS-I and LPS pathways. We also observed genetic mutation or transcriptional downregulation of other virulence genes in latter CF strains including TFP7 loci, Burkholderia lethal factor 1, T3SS-3 loci and flagellum loci, suggesting that these loci are similarly detrimental to long-term *B. pseudomallei* survival in the human host. Importantly, the RNA-seq data identified additional cases of surface antigen and virulence factor abrogation that were not observable with only genomic data. This finding underscores the importance of using both genomic and transcriptomic approaches to identify the functional consequences of within-host evolution of chronic bacterial infections.

TMP/SMX is used during the eradication phase of melioidosis treatment and is recommended for post-exposure prophylaxis (Peacock et al. 2008). We have previously shown that the latter isolate from CF6, and both isolates from CF11, had developed high-level (≥32 μg/mL) TMP/SMX resistance over the course of treatment. These elevated MICs were proposed to be due to mutations within BpeEF-OprC efflux pump regulators (BpeT T314fs in CF6, and BpeS V40I and R247L in CF11) alongside mutations affecting the R20 residue of Ptr1/FolM (R20fs in CF6; R20-A22 duplication in CF11) (Viberg et al. 2017). Here, we have demonstrated that the efflux pump regulatory mutations cause a dramatic upregulation of *bpeEF-oprC* in these strains of between 5.2- and 6.9-fold (38 to 121x), mirroring expression levels in *bpeS* and *bpeT* lab-generated mutants with high-level TMP/SMX resistance (Podnecky et al. 2017). Our results confirm those of Podnecky and colleagues (Podnecky et al. 2017) showing that upregulation of *bpeEF-oprC* via BpeS or BpeT dysregulation, together with Ptr1/FolM alteration, leads to a significant increase in TMP/SMX MICs that would render this antibiotic ineffective *in vivo*. RNA-seq is thus a useful tool for confirming the functional consequences of regulatory mutations that control RND efflux pump expression.

In addition to TMP/SMX resistance, the initial isolate from CF11 is resistant to CAZ (12 μg/mL) and the latter isolate from CF6 is highly resistant to CAZ (≥256 μg/mL). Our prior genomic study showed that CAZ resistance in the initial CF11 strain was due to a 10x duplication of a 36.7kb region encompassing the β-lactamase gene, *penA*, the first time that gene duplication has been shown to confer CAZ resistance in *B. pseudomallei* (Viberg et al. 2017). In contrast, a 2x duplication of this region in the latter strain did not raise the CAZ MIC above wild-type levels (Viberg et al. 2017). Similarly, the latter strain from CF6 exhibited a 30x duplication of a 7.5kb region encompassing *penA;* however, all 30 copies encoded a C69Y missense mutation, which by itself causes high-level (≥256 μg/mL) CAZ MICs (Sam et al. 2009). As anticipated, RNA-seq provided confirmation of the effects of gene duplications affecting *penA*, with the 30x duplication event in the latter CF6 strain resulting in a 4.5-fold (22x) corresponding increase in DE of this gene. Similarly, a 2.1-fold (4x) downregulation of *penA* in the latter strain from CF11 was linked to a 5x greater copy number of this gene in the early strain (Viberg et al. 2017). Thus, RNA-seq provided excellent correlation with gene copy number variation determined from whole-genome sequence coverage data. Taken together, the combined genetic and transcriptional changes affecting antibiotic resistance genes in the CF airways-adapted *B. pseudomallei* strains illustrates both the intractability of eradicating chronic bacterial infections and the unintended consequences of prolonged antibiotic use in CF treatment.

Although determining within-host transcriptional differences in longitudinal isolates yields valuable insights into the infection dynamics within individual patients, identifying convergent transcriptional changes provides a potential means to predict pathogen behavior and evolution across multiple CF cases in a relatively straightforward manner. Such predictability could conceivably be exploited to improve the diagnosis or treatment of intractable CF infections, or ideally, to prevent them from progressing in the first place. Therefore, a major objective of this study was to identify evidence of convergence in *B. pseudomallei* gene expression during its transition to a chronic infection. Despite the small number of CF melioidosis patients available for this study, a signal of convergent pathoadaptation was identified between the initial and latter isolates, with 19 significantly DE loci identified, 17 of which were upregulated (Table S2). This convergence is noteworthy given the large size of the *B. pseudomallei* genome and the many redundant pathways that could lead to similar adaptive phenotypes, a phenomenon that is well-recognized in *P. aeruginosa* (Marvig et al. 2015). One advantage of identifying convergence using transcriptomics rather than genomic data is that it can reveal the transcriptional consequence of multiple genetic mutations; for example, we have observed that multiple missense mutations in the RND efflux pump regulator AmrR lead to the same transcriptional outcome of *amrAB-oprA* upregulation (Sarovich et al. 2017). As such, RNA-seq data can simplify the identification of convergently expressed loci that are under the influence of several genetic variants.

The development of antibiotic resistance is a recurring theme in *P. aeruginosa* isolated from the CF airways (Winstanley et al. 2016), and we have recently shown that the same adaptive phenomenon can be observed in the genome of *B. pseudomallei* in response to prolonged, high-dose antibiotic therapy (Viberg et al. 2017). It was therefore not surprising to identify the convergent upregulation of *bpeEF-oprC* (4.8- to 6.1-fold; 28 to 69x), which was significantly upregulated in two of the four patients with DE (CF6 and CF11), and which led to TMP/SMX resistance as discussed above. The second convergently upregulated locus was the *cydAB* operon (*BPSL0501* and *BPSL0502*), which encodes for cytochrome *bd* quinol oxidase (5.5- to 5.9-fold; 45 to 60x); this locus was significantly DE in CF6 and CF9. CydAB is an aerobic terminal oxidase that oxidizes ubiquinol-8 and reduces oxygen to water under oxygen-limiting conditions. This enzyme is better able to scavenge oxygen under microaerobic conditions compared with cytochrome *o* oxidase, which otherwise predominates as the terminal respiratory enzyme in electron transport-associated energy production (Cotter et al. 1997). Voggu and colleagues demonstrated that the *cydAB* loci encoded by non-pathogenic *Staphylococcus* species were better able to resist *P. aeruginosa* antagonism in the CF lung compared with the *cydAB* loci encoded by *S. aureus*. This resistance was imparted by an insensitivity of the non-pathogenic staphylococci cytochrome *bd* quinol oxidases to the presence of the small respiratory inhibitors hydrogen cyanide and pyocyanin, which are commonly secreted by *P. aeruginosa* in the CF lung. In contrast, Voggu *et al*. showed that *S. aureus* was exquisitely sensitive to the co-presence of *P. aeruginosa* due to their less resistant *cydB* locus, which is inhibited by these small respiratory inhibitors (Voggu et al. 2006). Thus, it is feasible that the convergent upregulation of *cydAB* loci represents a defense mechanism employed by *B. pseudomallei* to counteract the toxic effects of small respiratory inhibitors produced by *P. aeruginosa* in the CF lung. In support of this hypothesis, *P. aeruginosa* was co-isolated in all five CF cases examined in this study (Viberg et al. 2017). Alternatively, *cydAB* upregulation may simply represent a physiological response to the oxygen-limited environment of the CF airways, as its expression is known to be induced in *B. pseudomallei* in hypoxic conditions (Hamad et al. 2011). Under such conditions, many pathogens including *B. pseudomallei* become less susceptible to conventional antibiotics, which are typically more effective under aerobic conditions, but more susceptible to antibiotics that target anaerobic infections, such as the nitroimidazole class of antibiotics (Hamad et al. 2011). This phenomenon may explain the difficulty of chronic *B. pseudomallei* eradication using conventional antibiotics like CAZ and TMP/SMX, and raises the exciting but not yet tested possibility that nitroimidazoles may be a highly effective therapeutic option for chronic, hypoxia-adapted *B. pseudomallei* infections such as those adapted to the CF airways.

A third convergently upregulated locus, the quorum sensing operon *hhqABCDEFG* (3.4- to 4.1-fold; 11x to 17x), is homologous to the *B. cepacia* complex *hmqABCDEFG* operon (Chapalain et al. 2017). This operon synthesizes a class of compounds known as 4-hydroxy-3-methyl-2-alkylquinolines (HMAQs), the methylated counterparts of 2-alkyl-4(1H)-quinolones (AHQs; also known as HAQs). AHQs were first recognized in *P. aeruginosa* and are produced by the signaling system *pqsABCDE* (Diggle et al. 2006). This cluster produces over 50 different AHQs, and these compounds exhibit diverse biological activities that enable cell-to-cell communication within and between bacterial species and the regulation of various functions including secondary metabolism, virulence, antibacterial activity and biofilm formation (Diggle et al. 2006). In contrast, little is currently known about the role of HMAQs and AHQs in *Burkholderia* spp. (Chapalain et al. 2017). The AHQ precursor molecule 2-heptyl-4(1H)-quinolone (HHQ) that is produced by *P. aeruginosa* actively suppresses the host innate immune response (Kim et al. 2010), a role that could be shared by *B. pseudomallei* HHQ. A second possibility is that these compounds impart a competitive advantage in the CF lung environment as HMAQs produced by *B. cepacia* exhibit antifungal activity (Kilani-Feki et al. 2011), so it is feasible that the *hmqABCDEFG* operon of *B. pseudomallei* produces similarly potent compounds that can inhibit fungal species from establishing residence in the CF lung. The convergent upregulation of *hhqA-G* in the *B. pseudomallei* CF isolates points to a putative role for AHQ-based compounds in *B. pseudomallei* signaling, immune evasion or competition in the CF lung. More work is needed to elucidate the myriad functions of AHQ compounds in *B. pseudomallei*, and particularly their role in promoting bacterial persistence in the CF airways.

Of the two convergently downregulated loci, only one, *BPSS0351*, has an assigned function, although little is known about the role of this gene and its product in *B. pseudomallei*. This gene encodes CueR (3.4-fold; 11x), a MerR family copper response regulator that is highly sensitive to the presence of copper (Cu) and which regulates the transcription of genes that protect against toxic metal ion concentrations (Brown et al. 2003; Singh et al. 2004). Cu has a long history as an effective antimicrobial agent due its ability to generate reactive oxygen species, with Cu accumulation in the mammalian host purported to act as an innate immune defense mechanism to restrict pathogen growth (Samanovic et al. 2012). Thus, downregulation of *cueR* in the latter CF isolates may represent a mechanism for mitigating Cu toxicity in the host, similarly to *E. coli* (Singh et al. 2004). However, there are contradictory reports as to whether Cu levels are elevated in CF sputa (Gray et al. 2010; Smith et al. 2014), and the artificial sputum growth medium does not appear to contain elevated Cu levels (Fung et al. 2010). CueR regulates the Cu/silver ATPase CopA and the multicopper oxidase CueO enzymes in *E. coli*, which correspond to *BPSS0224* and *BPSL0897* in *B. pseudomallei* K96243, respectively; however, neither of these genes were DE in any of the patient pairs. The absence of concomitant increased expression of *cueO*, which converts periplasmic Cu+ to less toxic Cu2+ *in vivo* (Singh et al. 2013), suggests that other enigmatic pressures are responsible for decreasing *cueR* expression in latter CF isolates. The biological role of these other factors requires further exploration.

We recognize that there are limitations to our study. Growth conditions are known to be an important consideration for mRNA-based investigations due to the alteration of the transcriptome when isolates are grown under different environments or media components (Yoder-Himes et al. 2009). Although our *in vitro* conditions do not completely mimic the conditions seen in the CF lung, the artificial sputum medium is designed to reflect the nutrient conditions of this environment (Fung et al. 2010), and our shaking parameters provided a robust way of measuring cellular growth over time while avoiding non-uniform cellular growth, which ensured harvest of *B. pseudomallei* cultures at the same growth phase (Figure S1). Additionally, the use of isogenic strain pairs with genomic data (Viberg et al. 2017) enabled us to comprehensively assess the effects of transcriptional adaptation to the CF lung compared with their underlying genetic variants. Our conditions provided transcriptomic data that was consistent with expected expression differences based on genome-wide alterations. Other studies have used artificial sputum media and additional mechanical methods to mimic the CF lung conditions. A rotating wall vessel has been developed to simulate the low fluid shear conditions encountered in CF mucus due to pathological effects of *CFTR* dysfunction on mucociliary clearance (Crabbé et al. 2008), with CF-derived *P. aeruginosa* isolates demonstrating transcriptional differences depending on shear conditions (Dingemans et al. 2016) . The culturing methods for bacterial RNA-seq are a critical consideration in experimental design as they can affect transcriptomic profiles, and the impact of conditions should be considered when comparing transcriptional differences between studies. Another shortcoming is that we only examined five patient pairs due to the relative paucity of melioidosis CF cases worldwide, and only two isolates from each patient due to limited bacterial colony selection and storage at the time of sputum collection and processing. Deeper sampling efforts across a greater number of melioidosis CF patients would be needed to provide greater confidence in our convergent adaptation findings and would allow a more advanced and detailed understanding of *B. pseudomallei* population dynamics and diversity to be attained. Nevertheless, the findings from our study provide important new insights into *B. pseudomallei* evolution in the CF airways, with many, although not complete, parallels with the common CF pathogens, *P. aeruginosa* and *B. cepacia* complex species.

## METHODS

### Ethics statement

Ethics approval for this study was obtained as previously described (Currie et al. 2010; Geake et al. 2015).

### CF isolates

The *B. pseudomallei* strains used in this study are summarized in Table 1. The history and genomic analysis of these cases and strains are detailed elsewhere (Viberg et al. 2017).

### Artificial sputum medium

This medium was made as previously described (Sriramulu et al. 2005; Fung et al. 2010), with modifications detailed here. Antibiotics were not used to maintain media sterility due to concerns that their addition would alter expression profiles. Due to impracticality in its filtration (Dingemans et al. 2016), 1 g porcine stomach mucin (Sigma-Aldrich, Castle Hill, NSW, Australia) dissolved in 40 mL ultrapure water was autoclaved prior to use. All other solutions were sterilized using a 0.22 μM vacuum filter, apart from the UV-irradiated egg yolk emulsion (Oxoid, Thebarton, SA, Australia), which was treated aseptically. A stock solution of diethylenetriaminepentaacetic acid (Sigma-Aldrich) was made by dissolving 59.5 mg into 5 mL of very basic water (pH=14). CaCl_2_ was added at a final concentration of 0.22 g/L (J. Manos, pers. comm.). Final concentrations of the components were: 10 g/L mucin, 1.39 g/L salmon sperm DNA (Sigma-Aldrich), 5 g/L NaCl, 2.2 g/L KCl, 0.22 g/L CaCl_2_, 5 g/L casein acid hydrosylate (Sigma-Aldrich), 10 g/L bovine serum albumin (Roche Diagnostics, Castle Hill, NSW, Australia), 0.005% diethylenetriaminepentaacetic acid, and 0.5% of egg yolk emulsion. Each batch was tested for sterility prior to use by plating 100 μL onto Luria Bertani (LB) agar (Oxoid) and incubating aerobically for 24 h. pH was tested using an aliquot of the medium to ensure it was within the desired range (pH ~6.5-7). The medium was stored at 4 ^o^C for no longer than four weeks prior to use.

### Viability counts

Two sets of viability counts were performed for this study. The first was conducted to determine the number of colony-forming units (cfu) at OD_590_, which enabled us to standardize the starting number of cells inoculated into the artificial sputum medium. The second was conducted to verify the final concentration of cells across all CF isolates, which enabled us to determine the number of cells for nucleic acid extraction to ensure that approximately equal cell amounts were processed for each pair. The CF isolates were subcultured from glycerol stocks onto LB agar at 37 °C for 24 h. Cells were suspended into phosphate-buffered saline (PBS) followed by spectrophotometric measurement at OD_590_=1.0 in a WPA CO 8000 cell density meter (Biochrom Ltd, Blackburn, VIC, Australia). Tenfold dilutions and plating of cultures onto LB agar was carried out, followed by enumeration at 24 h. Viable counts demonstrated that all CF isolates exhibited similar cell density when normalized to an OD_590_=1.0 (range 1.3×10^8^ to 4.9×10^8^). Based on these counts, the starting amount of culture for the CF isolates was standardized to 10^5^ cfu for all subsequent experiments.

### Growth curves in the artificial sputum medium

To minimize laboratory passage, each culture was again subcultured from the original glycerol stocks onto LB agar at 37 °C for 24 h, followed by a replication of OD_590_ measurements as determined previously. Based on the viability count data, samples were then diluted to 10^6^ cfu/mL in PBS. One hundred μL of this suspension (~10^5^ cfu) was used to inoculate 1.9 mL of the sputum medium, which was aliquoted into 14 mL Nunc round-bottom culture tubes (Thermo Fisher Scientific, Scoresby, VIC, Australia). Due to biosafety concerns, cells were grown in closed-capped tubes, and were incubated at 37 °C by shaking in an orbital incubator shaker (model BL8500; Bioline, Eveleigh, NSW, Australia) at 50, 200 or 230 rpm for 44 h. Growth curves were obtained by measuring OD_590_ at regular intervals over this period using un-inoculated sputum medium as the control blank. Shaking at 50 rpm was initially performed to mimic the low-oxygen conditions of the CF lung; however, this speed caused heavy sedimentation of cells and media components, and biofilm formation at the aerobic interface, both of which led to a decrease in OD values over time and unpredictable, non-uniform growth. Similarly, shaking at 230 rpm was too vigorous for the cells, as observed by inconsistent, non-reproducible viable counts. When shaking at 200 rpm, highly reproducible OD values that correlated with viability counts were obtained (Figure S1), and the medium did not readily sediment. We therefore used this speed for subsequent experiments, including for RNA harvest. Viability counts were performed at the time of harvest to ensure uniformity of cell concentrations across all isolates.

### RNA extraction from isolates grown in the artificial sputum medium

The CF strains were grown in duplicate at 200 rpm as detailed above. Based on the growth curve analysis, nucleic acids for all cultures were extracted at late log phase (17 h). At the point of harvest, the OD_590_ of each replicate was measured to ensure that consistent cell density had been obtained prior to combining replicates; final viability counts were also performed. Due to the highly labile nature of bacterial mRNA, two 100 μL aliquots for each strain were immediately placed into 200 μL of RNAprotect (Qiagen, Doncaster, VIC, Australia) and incubated for 5 min to preserve their transcription profiles. Cells were pelleted by centrifugation at 5000 x *g* for 10 min, and the supernatant discarded. Total RNA was extracted using the RNeasy Protect Bacteria Mini Kit (Qiagen). *B. pseudomallei* cells were lysed following the protocol for genomic DNA extraction (Currie et al. 2007), with an extended incubation time in Proteinase K to 1.5 h. Lysates were loaded onto the RNeasy mini columns and extractions were carried out according to the manufacturer’s instructions, including the recommended on-column DNase I digestion. In our hands, we found this DNase I treatment to be insufficient for removing all contaminating DNA. Extractions for RNA-seq were therefore further treated with TURBO DNA-*free* kit (Ambion, Scoresby, VIC, Australia). For each sample, 35 μL of extracted RNA was incubated with 6 U TURBO DNase at 37 °C for 32 min. The remaining RNA was not treated with this second round of DNase; instead, this sample was used as template for PCR contamination screening, as described below. All samples were transferred to clean RNase/DNase-free tubes for downstream processing.

### RNA quality control

To verify the removal of DNA from the total RNA extractions, two contamination screens were performed. The first was used to determine the removal of salmon sperm DNA, and the second was to determine the removal of *B. pseudomallei* DNA. Both the pre- and post-treated RNA samples were used to test for contamination, in duplicate, with the former acting as the positive control. The RNA samples were diluted 1/10 into molecular-grade H2O (Fisher Biotech) prior to PCR. Identification of residual salmon sperm DNA was investigated by targeting the mitochondrial 12S rDNA region of vertebrates (Humair et al. 2007). Primers 12S-6F (5’- CAAACTGGGATTAGATACC) and B-12S-9R (5’- AGAACAGGCTCCTCTAG) were used at a final concentration of 1 μM in a mix containing 1X PCR buffer (Qiagen), 1 U HotStarTaq, 0.2 mM dNTPs, 1 μL template and molecular-grade H2O in a 15 μL total reaction volume. Thermocycling conditions comprised 94 °C for 5 min, followed by 35 cycles of 94 °C for 30 sec, 52 °C for 30 sec, and 72 °C for 30 sec, and a final extension at 72 °C for 2 min. Amplicons were detected by agarose gel electrophoresis.

Real-time PCR was used to detect *B. pseudomallei* DNA contamination. The *mmsA* (methylmalonate-semialdehyde dehydrogenase) housekeeping gene was targeted using the primers Bp_266152_3012-F1-flap (5’-AATAAATCATAAACGTGAGGCCGGAGATGT) and Bp_266152_3012-R1-flap (5’-AATAAATCATAAGACCGACATCACGCACAGC) in combination with a *B. pseudomallei-specific* TaqMan MGB probe, 266152-T_Bp (5’-VIC-CGGTCTACACGCATGA) (Price et al. 2012), as previously described, with the following modifications: 0.2 μM probe and 0.4 μM each primer was used, reactions were carried out in a 5 μL total reaction volume, and cycling was performed to 50 cycles.

### RNA storage, shipment and RNA-seq

For each sample, 20 μL of total RNA was added to an RNAstable tube (Biomatrica, San Diego, CA, USA), gently mixed with the preservation agent and left to air-dry in a biosafety cabinet for 48 h. Samples were shipped at ambient temperature to Macrogen Inc. (Geumcheon-gu, Seoul, Rep. of Korea) for RNA-seq. Ribosomal RNA was removed by treatment with the Ribo-Zero rRNA Removal Kit for Bacteria (Epicentre, Madison, WI, USA), followed by 100 bp paired-end, stranded library construction using the TruSeq rapid SBS Kit (Illumina Inc., San Diego, CA, USA). Libraries were sequenced on either the HiSeq2000 or HiSeq2500 platform (Illumina Inc.). All samples were extracted from two separate experiments to account for biological variation, except for MSHR8442, which was extracted thrice. Between 36 and 80 million reads were generated for each sequence, corresponding to between 3.6 and 8.1 billion base pairs each.

### RNA-seq analysis

Illumina read filtering was first performed with Trimmomatic v0.33 using the following parameters: TruSeq2-PE adapter removal, leading=3, trailing=3, sliding window=4:15, and minimum length=36. Reads were mapped to the prototypic *B. pseudomallei* K96243 reference genome (RefSeq IDs NC_006350 and NC_006351 for chromosomes 1 and 2, respectively (Holden et al. 2004)) using Bowtie 2 v2.2.1 (Langmead and Salzberg 2012). Transcript quantification was performed with HTSeq (v0.6.1p1) (Anders et al. 2015) using the intersection non-empty mode and --stranded=reverse parameters. DE analysis was carried out using the glmFit function of edgeR v3.18.1 (Robinson et al. 2010), implemented in the online Degust 3.1.0 tool (http://www.vicbioinformatics.com/degust/index.html). DE loci were visualized using the volcano plot function within Degust. Several different groups were compared to determine DE. The first analyses compared initial and latter isolates within CF patients (Table 1) without summing technical replicates (i.e. the RNA-seq data from each independent experiment of a single strain) to identify DE within each patient. To determine convergent DE loci, we summed the reads for each technical replicate prior to analysis and then compared all initial CF isolates vs. all latter CF isolates, with the latter CF10 isolate excluded due to a lack of DE in this strain and the initial CF11 isolate excluded due to >3 years of infection prior to its isolation (Viberg et al. 2017). For all analyses, DE was defined as a log2 fold change of ≥1.5 and a false discovery rate of ≤0.01. To improve visualization of DE loci in the volcano plot of the initial and latter comparison (Figure 1), highly expressed DE loci in only a single strain were omitted.

## DATA ACCESS

The RNA-seq data generated in this study are available on the Sequence Read Archive database under BioProject number PRJNA398168 and submission numbers SRR6031143 to SRR6031152.

## ACKNOWLEDGEM ENTS

We are grateful to Dr Jim Manos (University of Sydney, NSW, Australia) for providing advice on the preparation of the artificial sputum medium, Ammar Aziz for helpful conversations regarding RNA-seq analyses, and Jessica Webb, Mark Mayo and Vanessa Theobald (Menzies School of Health Research) for laboratory assistance. This work was supported by grants 1046812 and 1098337 from the Australian National Health and Medical Research Council. The funder had no role in study design, data collection and interpretation, or the decision to submit the work for publication. EPP was supported by a University of the Sunshine Coast Fellowship, LTV was supported by an Australian Postgraduate Award and Menzies Enhanced Living scholarship, SCB was supported by a Health Research Fellowship from Queensland Health, and DSS was supported by an Advance Queensland Fellowship (award number AQRF13016-17RD2).

## AUTHOR CONTRIBUTIONS

EPP, BJC and DSS conceived of and obtained funding for the study. EPP, LTV and DSS designed the laboratory experiments LTV carried out experiments. EPP, LTV and DSS performed bioinformatic analyses. TJK, SCB and BJC supplied the *B. pseudomallei* isolates and clinical information. EPP and DSS wrote the manuscript. All authors read and approved the final manuscript.

## REFERENCES

Amaral MD. 2015. Novel personalized therapies for cystic fibrosis: treating the basic defect in all patients. J Intern Med 277: 155–166.

Ambler RP, Coulson AF, Frere JM, Ghuysen JM, Joris B, Forsman M, Levesque RC, Tiraby G, Waley SG. 1991. A standard numbering scheme for the class A beta-lactamases. Biochem J 276 (Pt 1): 269–270.

Anders S, Pyl PT, Huber W. 2015. HTSeq—a Python framework to work with high-throughput sequencing data. Bioinformatics 31: 166–169.

Atkins T, Prior R, Mack K, Russell P, Nelson M, Prior J, Ellis J, Oyston PC, Dougan G, Titball RW. 2002. Characterisation of an acapsular mutant of *Burkholderia pseudomallei* identified by signature tagged mutagenesis. J Med Microbiol 51: 539–547.

Beach MB, Osuna R. 1998. Identification and characterization of the *fis* operon in enteric bacteria. J Bacteriol 180: 5932–5946.

Brown NL, Stoyanov JV, Kidd SP, Hobman JL. 2003. The MerR family of transcriptional regulators. FEMS Microbiol Rev 27: 145–163.

Chapalain A, Groleau MC, Le Guillouzer S, Miomandre A, Vial L, Milot S, Deziel E. 2017. Interplay between 4-hydroxy-3-methyl-2-alkylquinoline and N-Acyl-homoserine lactone signaling in a *Burkholderia cepacia* complex clinical strain. Front Microbiol 8: 1021.

Chapple SN, Sarovich DS, Holden MT, Peacock SJ, Buller N, Golledge C, Mayo M, Currie BJ, Price EP. 2016. Whole-genome sequencing of a quarter-century melioidosis outbreak in temperate Australia uncovers a region of low-prevalence endemicity. Microb Genom 2: e000067.

Cohen TS, Prince A. 2012. Cystic fibrosis: a mucosal immunodeficiency syndrome. Nat Med 18: 509–519.

Cotter PA, Melville SB, Albrecht JA, Gunsalus RP. 1997. Aerobic regulation of cytochrome *d* oxidase *(cydAB)* operon expression in *Escherichia coli:* roles of Fnr and ArcA in repression and activation. Mol Microbiol 25: 605–615.

Coutinho HD, Falcao-Silva VS, Goncalves GF. 2008. Pulmonary bacterial pathogens in cystic fibrosis patients and antibiotic therapy: a tool for the health workers. Int Arch Med 1: 24.

Crabbé A, De Boever P, Van Houdt R, Moors H, Mergeay M, Cornelis P. 2008. Use of the rotating wall vessel technology to study the effect of shear stress on growth behaviour of *Pseudomonas aeruginosa* PA01. Environ Microbiol 10: 2098–2110.

Creecy JP, Conway T. 2015. Quantitative bacterial transcriptomics with RNA-seq. Curr Opin Microbiol 23: 133–140.

Cuccui J, Milne TS, Harmer N, George AJ, Harding SV, Dean RE, Scott AE, Sarkar-Tyson M, Wren BW, Titball RW et al. 2012. Characterization of the *Burkholderia pseudomallei* K96243 capsular polysaccharide I coding region. Infect Immun 80: 1209–1221.

Currie BJ. 2015. Melioidosis: evolving concepts in epidemiology, pathogenesis, and treatment. Semin Respir Crit Care Med 36: 111–125.

Currie BJ, Gal D, Mayo M, Ward L, Godoy D, Spratt BG, LiPuma JJ. 2007. Using BOX-PCR to exclude a clonal outbreak of melioidosis. BMC Infect Dis 7: 68.

Currie BJ, Ward L, Cheng AC. 2010. The epidemiology and clinical spectrum of melioidosis: 540 cases from the 20 year Darwin prospective study. PLoS Negl Trop Dis 4: e900.

DeShazer D, Brett PJ, Woods DE. 1998. The type II O-antigenic polysaccharide moiety of *Burkholderia pseudomallei* lipopolysaccharide is required for serum resistance and virulence. Mol Microbiol 30: 1081–1100.

DeShazer D, Waag DM, Fritz DL, Woods DE. 2001. Identification of a *Burkholderia mallei* polysaccharide gene cluster by subtractive hybridization and demonstration that the encoded capsule is an essential virulence determinant. Microb Pathog 30: 253–269.

Diggle SP, Lumjiaktase P, Dipilato F, Winzer K, Kunakorn M, Barrett DA, Chhabra SR, Camara M, Williams P. 2006. Functional genetic analysis reveals a 2-alkyl-4-quinolone signaling system in the human pathogen *Burkholderia pseudomallei* and related bacteria. Chem Biol 13: 701–710.

Dingemans J, Monsieurs P, Yu SH, Crabbe A, Forstner KU, Malfroot A, Cornelis P, Van Houdt R. 2016. Effect of shear stress on *Pseudomonas aeruginosa* isolated from the cystic fibrosis lung. MBio 7: e00813-16.

Dvorak GD, Spickler AR. 2008. Glanders. J Am Vet Med Assoc 233: 570–577.

Fung C, Naughton S, Turnbull L, Tingpej P, Rose B, Arthur J, Hu H, Harmer C, Harbour C, Hassett DJ et al. 2010. Gene expression of *Pseudomonas aeruginosa* in a mucin-containing synthetic growth medium mimicking cystic fibrosis lung sputum. J Med Microbiol 59: 1089–1100.

Geake JB, Reid DW, Currie BJ, Bell SC, Melioid CFI, Bright-Thomas R, Dewar J, Holden S, Simmonds N, Gyi K et al. 2015. An international, multicentre evaluation and description of *Burkholderia pseudomallei* infection in cystic fibrosis. BMC Pulm Med 15: 116.

Gray RD, Duncan A, Noble D, Imrie M, O’Reilly DS, Innes JA, Porteous DJ, Greening AP, Boyd AC. 2010. Sputum trace metals are biomarkers of inflammatory and suppurative lung disease. Chest 137: 635–641.

Hamad MA, Austin CR, Stewart AL, Higgins M, Vazquez-Torres A, Voskuil MI. 2011. Adaptation and antibiotic tolerance of anaerobic *Burkholderia pseudomallei*. Antimicrob Agents Chemother 55: 3313–3323.

Holden MT, Titball RW, Peacock SJ, Cerdeno-Tarraga AM, Atkins T, Crossman LC, Pitt T, Churcher C, Mungall K, Bentley SD et al. 2004. Genomic plasticity of the causative agent of melioidosis, *Burkholderia pseudomallei*. Proc Natl Acad Sci U S A 101: 14240–14245.

Holland DJ, Wesley A, Drinkovic D, Currie BJ. 2002. Cystic fibrosis and *Burkholderia pseudomallei* infection: an emerging problem? Clin Infect Dis 35: e138-140.

Humair PF, Douet V, Moran Cadenas F, Schouls LM, Van De Pol I, Gern L. 2007. Molecular identification of bloodmeal source in *Ixodes ricinus* ticks using 12S rDNA as a genetic marker. J Med Entomol 44: 869–880.

Kilani-Feki O, Culioli G, Ortalo-Magne A, Zouari N, Blache Y, Jaoua S. 2011. Environmental *Burkholderia cepacia* strain Cs5 acting by two analogous alkyl-quinolones and a didecyl-phthalate against a broad spectrum of phytopathogens fungi. Curr Microbiol 62: 1490–1495.

Kim K, Kim YU, Koh BH, Hwang SS, Kim SH, Lepine F, Cho YH, Lee GR. 2010. HHQ and PQS, two *Pseudomonas aeruginosa* quorum-sensing molecules, down-regulate the innate immune responses through the nuclear factor-kappaB pathway. Immunology 129: 578588.

Langmead B, Salzberg SL. 2012. Fast gapped-read alignment with Bowtie 2. Nat Methods 9: 357–359.

Leelarasamee A, Bovornkitti S. 1989. Melioidosis: review and update. Rev Infect Dis 11: 413425.

Lipsitz R, Garges S, Aurigemma R, Baccam P, Blaney DD, Cheng AC, Currie BJ, Dance D, Gee JE, Larsen J et al. 2012. Workshop on treatment of and postexposure prophylaxis for *Burkholderia pseudomallei* and *B. mallei* infection, 2010. Emerg Infect Dis 18: e2.

Marvig RL, Sommer LM, Jelsbak L, Molin S, Johansen HK. 2015. Evolutionary insight from whole-genome sequencing of *Pseudomonas aeruginosa* from cystic fibrosis patients. Future Microbiol 10: 599–611.

Moore RA, DeShazer D, Reckseidler S, Weissman A, Woods DE. 1999. Efflux-mediated aminoglycoside and macrolide resistance in *Burkholderia pseudomallei*. Antimicrob Agents Chemother 43: 465–470.

Nierman WC, DeShazer D, Kim HS, Tettelin H, Nelson KE, Feldblyum T, Ulrich RL, Ronning CM, Brinkac LM, Daugherty SC et al. 2004. Structural flexibility in the *Burkholderia mallei* genome. Proc Natl Acad Sci U S A 101: 14246–14251.

O’Carroll MR, Kidd TJ, Coulter C, Smith HV, Rose BR, Harbour C, Bell SC. 2003. *Burkholderia pseudomallei:* another emerging pathogen in cystic fibrosis. Thorax 58: 1087–1091.

Ooi WF, Ong C, Nandi T, Kreisberg JF, Chua HH, Sun G, Chen Y, Mueller C, Conejero L, Eshaghi M et al. 2013. The condition-dependent transcriptional landscape of *Burkholderia pseudomallei*. PLoS Genet 9: e1003795.

Peacock SJ, Schweizer HP, Dance DA, Smith TL, Gee JE, Wuthiekanun V, DeShazer D, Steinmetz I, Tan P, Currie BJ. 2008. Management of accidental laboratory exposure to *Burkholderia pseudomallei* and *B. mallei*. Emerg Infect Dis 14: e2.

Podnecky NL, Rhodes KA, Mima T, Drew HR, Chirakul S, Wuthiekanun V, Schupp JM, Sarovich DS, Currie BJ, Keim P et al. 2017. Mechanisms of resistance to folate pathway inhibitors in *Burkholderia pseudomallei:* deviation from the norm. MBio 8: e01357-17.

Price EP, Dale JL, Cook JM, Sarovich DS, Seymour ML, Ginther JL, Kaufman EL, Beckstrom-Sternberg SM, Mayo M, Kaestli M et al. 2012. Development and validation of *Burkholderia pseudomallei-specific* real-time PCR assays for clinical, environmental or forensic detection applications. PLoS One 7: e37723.

Price EP, Sarovich DS, Mayo M, Tuanyok A, Drees KP, Kaestli M, Beckstrom-Sternberg SM, Babic-Sternberg JS, Kidd TJ, Bell SC et al. 2013. Within-host evolution of *Burkholderia pseudomallei* over a twelve-year chronic carriage infection. MBio 4: e00388-13.

Price EP, Sarovich DS, Viberg L, Mayo M, Kaestli M, Tuanyok A, Foster JT, Keim P, Pearson T, Currie BJ. 2015. Whole-genome sequencing of *Burkholderia pseudomallei* isolates from an unusual melioidosis case identifies a polyclonal infection with the same multilocus sequence type. J Clin Microbiol 53: 282–286.

Reckseidler-Zenteno SL, DeVinney R, Woods DE. 2005. The capsular polysaccharide of *Burkholderia pseudomallei* contributes to survival in serum by reducing complement factor C3b deposition. Infect Immun 73: 1106–1115.

Reckseidler-Zenteno SL, Viteri DF, Moore R, Wong E, Tuanyok A, Woods DE. 2010. Characterization of the type III capsular polysaccharide produced by *Burkholderia pseudomallei*. J Med Microbiol 59: 1403–1414.

Reckseidler SL, DeShazer D, Sokol PA, Woods DE. 2001. Detection of bacterial virulence genes by subtractive hybridization: identification of capsular polysaccharide of *Burkholderia pseudomallei* as a major virulence determinant. Infect Immun 69: 34–44.

Rholl DA, Papp-Wallace KM, Tomaras AP, Vasil ML, Bonomo RA, Schweizer HP. 2011. Molecular investigations of PenA-mediated beta-lactam resistance in *Burkholderia pseudomallei*. Front Microbiol 2: 139.

Robinson MD, McCarthy DJ, Smyth GK. 2010. edgeR: a Bioconductor package for differential expression analysis of digital gene expression data. Bioinformatics 26: 139–140.

Sam IC, See KH, Puthucheary SD. 2009. Variations in ceftazidime and amoxicillin-clavulanate susceptibilities within a clonal infection of *Burkholderia pseudomallei*. J Clin Microbiol 47: 1556–1558.

Samanovic MI, Ding C, Thiele DJ, Darwin KH. 2012. Copper in microbial pathogenesis: meddling with the metal. Cell Host Microbe 11: 106–115.

Sarovich DS, Garin B, De Smet B, Kaestli M, Mayo M, Vandamme P, Jacobs J, Lompo P, Tahita MC, Tinto H et al. 2016. Phylogenomic analysis reveals an Asian origin for African *Burkholderia pseudomallei* and further supports melioidosis endemicity in Africa. mSphere 1: e00089-00015.

Sarovich DS, Price EP, Von Schulze AT, Cook JM, Mayo M, Watson LM, Richardson L, Seymour ML, Tuanyok A, Engelthaler DM et al. 2012. Characterization of ceftazidime resistance mechanisms in clinical isolates of *Burkholderia pseudomallei* from Australia. PLoS One 7: e30789.

Sarovich DS, Webb JR, Pitman MC, Viberg LT, Mayo M, Baird RW, Robson JM, Currie BJ, Price EP. 2017. Raising the stakes: loss of efflux pump regulation decreases meropenem susceptibility in *Burkholderia pseudomallei*. bioRxiv doi:https://doi.org/10.1101/205070.

Sharma CM, Hoffmann S, Darfeuille F, Reignier J, Findeiss S, Sittka A, Chabas S, Reiche K, Hackermuller J, Reinhardt R et al. 2010. The primary transcriptome of the major human pathogen *Helicobacter pylori*. Nature 464: 250–255.

Singh AP, Lai SC, Nandi T, Chua HH, Ooi WF, Ong C, Boyce JD, Adler B, Devenish RJ, Tan P. 2013. Evolutionary analysis of *Burkholderia pseudomallei* identifies putative novel virulence genes, including a microbial regulator of host cell autophagy. J Bacteriol 195: 5487–5498.

Singh SK, Grass G, Rensing C, Montfort WR. 2004. Cuprous oxidase activity of CueO from *Escherichia coli*. J Bacteriol 186: 7815–7817.

Smith DJ, Anderson GJ, Bell SC, Reid DW. 2014. Elevated metal concentrations in the CF airway correlate with cellular injury and disease severity. J Cyst Fibros 13: 289–295.

Sriramulu DD, Lunsdorf H, Lam JS, Romling U. 2005. Microcolony formation: a novel biofilm model of *Pseudomonas aeruginosa* for the cystic fibrosis lung. J Med Microbiol 54: 667676.

Stone JK, DeShazer D, Brett PJ, Burtnick MN. 2014. Melioidosis: molecular aspects of pathogenesis. Expert Rev Anti Infect Ther 12: 1487–1499.

Tuanyok A, Stone JK, Mayo M, Kaestli M, Gruendike J, Georgia S, Warrington S, Mullins T, Allender CJ, Wagner DM et al. 2012. The genetic and molecular basis of O-antigenic diversity in *Burkholderia pseudomallei* lipopolysaccharide. PLoS Negl Trop Dis 6: e1453.

Viberg LT, Sarovich DS, Kidd TJ, Geake JB, Bell SC, Currie BJ, Price EP. 2017. Within-host evolution of *Burkholderia pseudomallei* during chronic infection of seven Australasian cystic fibrosis patients. MBio 8: e00356-17.

Voggu L, Schlag S, Biswas R, Rosenstein R, Rausch C, Gotz F. 2006. Microevolution of cytochrome *bd* oxidase in Staphylococci and its implication in resistance to respiratory toxins released by *Pseudomonas*. J Bacteriol 188: 8079–8086.

Webb JR, Price EP, Currie BJ, Sarovich DS. 2017. Loss of methyltransferase function and increased efflux activity leads to doxycycline resistance in *Burkholderia pseudomallei*. Antimicrob Agents Chemother 61 : e00268-00217.

Wiersinga WJ, Currie BJ, Peacock SJ. 2012. Melioidosis. N Engl J Med 367: 1035–1044.

Winstanley C, O’Brien S, Brockhurst MA. 2016. *Pseudomonas aeruginosa* evolutionary adaptation and diversification in cystic fibrosis chronic lung infections. Trends Microbiol 24: 327–337.

Yip TW, Hewagama S, Mayo M, Price EP, Sarovich DS, Bastian I, Baird RW, Spratt BG, Currie BJ. 2015. Endemic melioidosis in residents of desert region after atypically intense rainfall in central Australia, 2011. Emerg Infect Dis 21: 1038–1040.

Yoder-Himes DR, Chain PS, Zhu Y, Wurtzel O, Rubin EM, Tiedje JM, Sorek R. 2009. Mapping the *Burkholderia cenocepacia* niche response via high-throughput sequencing. Proc Natl Acad Sci U S A 106: 3976–3981.

